## Supplementary material for "Cross-feeding affects the target of resistance evolution to an antifungal drug": Supp Figs 1-10

### Supporting information

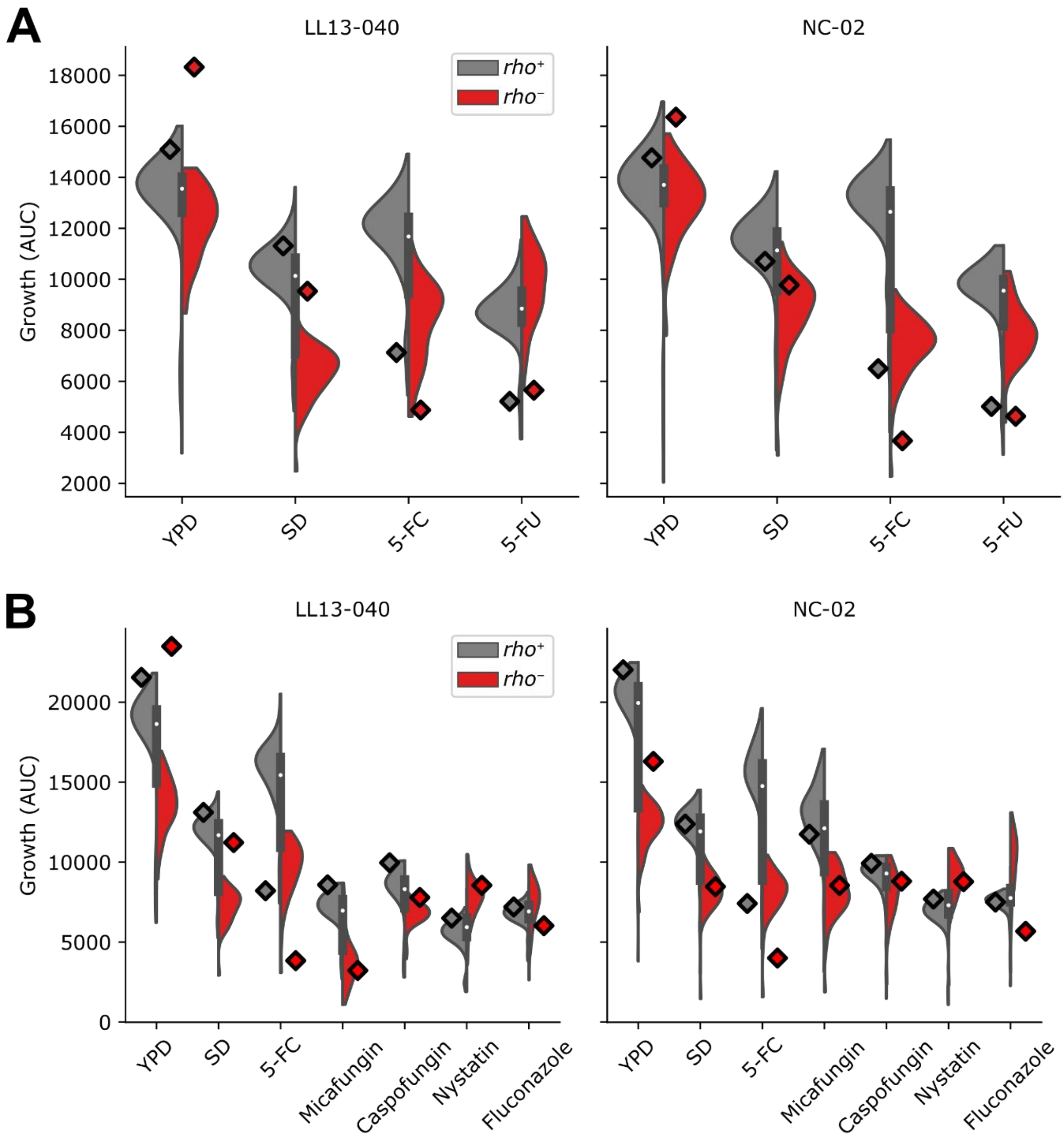


**Supp Fig 1. Growth of individual mutants on solid medium.** Growth corresponds to the mean area under the curve (AUC, calculated on 22 h) from four replicate colonies, for individual strains arrayed on solid media: (A) YPD, SD, SD + 25 µg/mL 5-FC and SD + 6.25 µg/mL 5-FU, incubated at 30°C and (B) YPD, SD, SD + 25 µg/mL 5-FC, SD + 0.5 µg/mL micafungin, SD + 2 µg/mL caspofungin, SD + 16 µg/mL nystatin and SD + 64 µg/mL fluconazole, incubated at 37°C. Data correspond to Figs 2 and 3A before normalization with the WT, here represented as split violin plots. For both growth assays (corresponding to panels A and B), all *rho*$^{+}$ strains were gathered on a single plate, with WT controls for the two backgrounds (LL13-040 and NC-02). The mean AUC for the WT controls in each condition is indicated by a grey diamond. Similarly, the mean AUC for the WT controls present on plates with *rho*$^{-}$ strains (in this case, one plate for LL13-040 *rho*$^{-}$ strains and another for NC-02 *rho*$^{-}$ strains) is indicated by a red diamond.

**Supp Fig 2. Depth per position.** Normalized coverage calculated for 5 kb windows. Each track corresponds to a sequenced genome. Red signal indicates a normalized coverage above 4X. First page corresponds to LL13-040 genomes (B10, B11…) and second page to NC-02 genomes (B14, B15…).

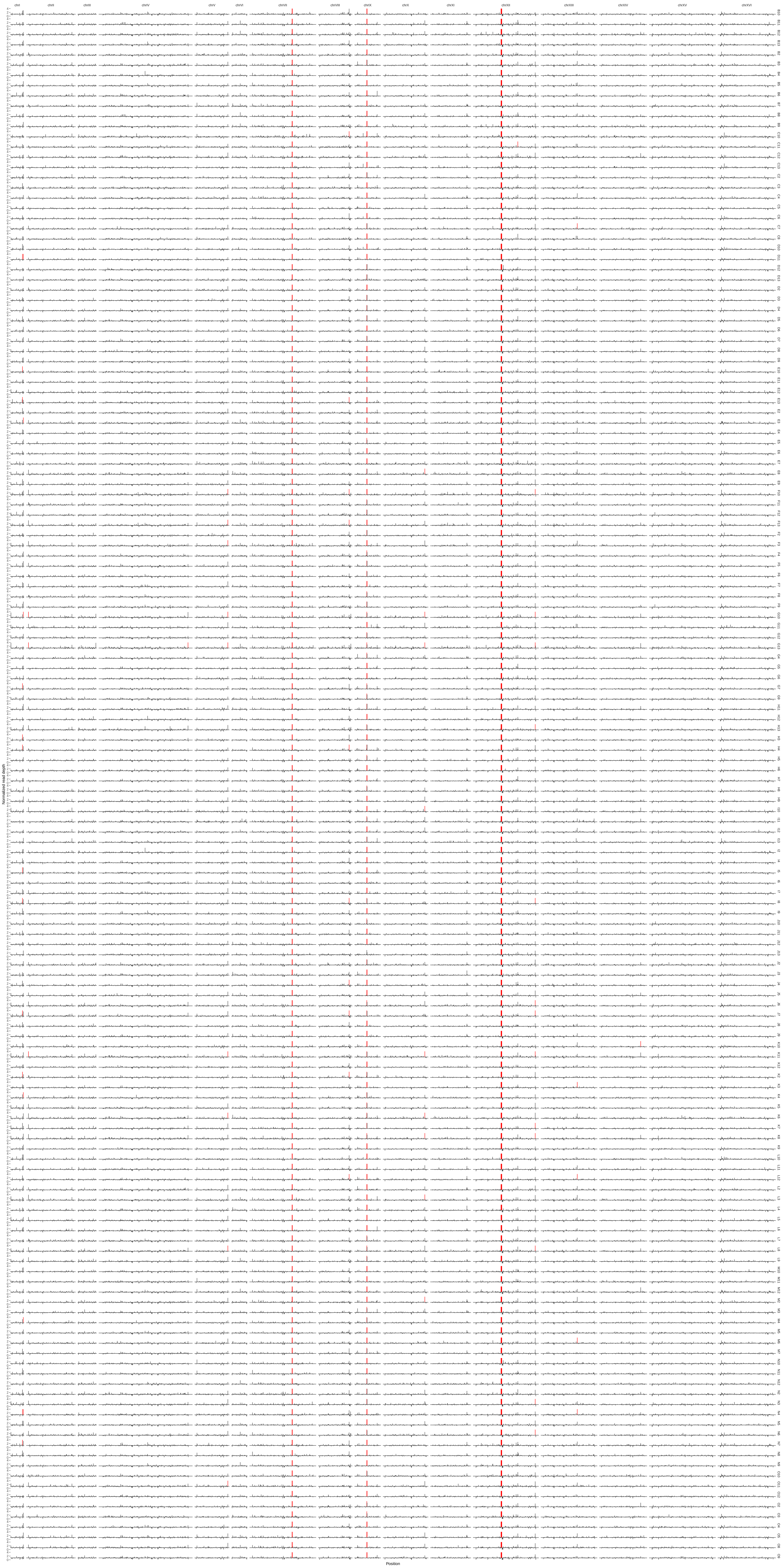
chrI
chrII
chrIII
chrIV
chrV
chrVI
chrVII
chrVIII
chrIX
chrX
chrXI
chrXII
chrXIII
chrXIV
chrXV
chrXVI
Normalized read depth
Position

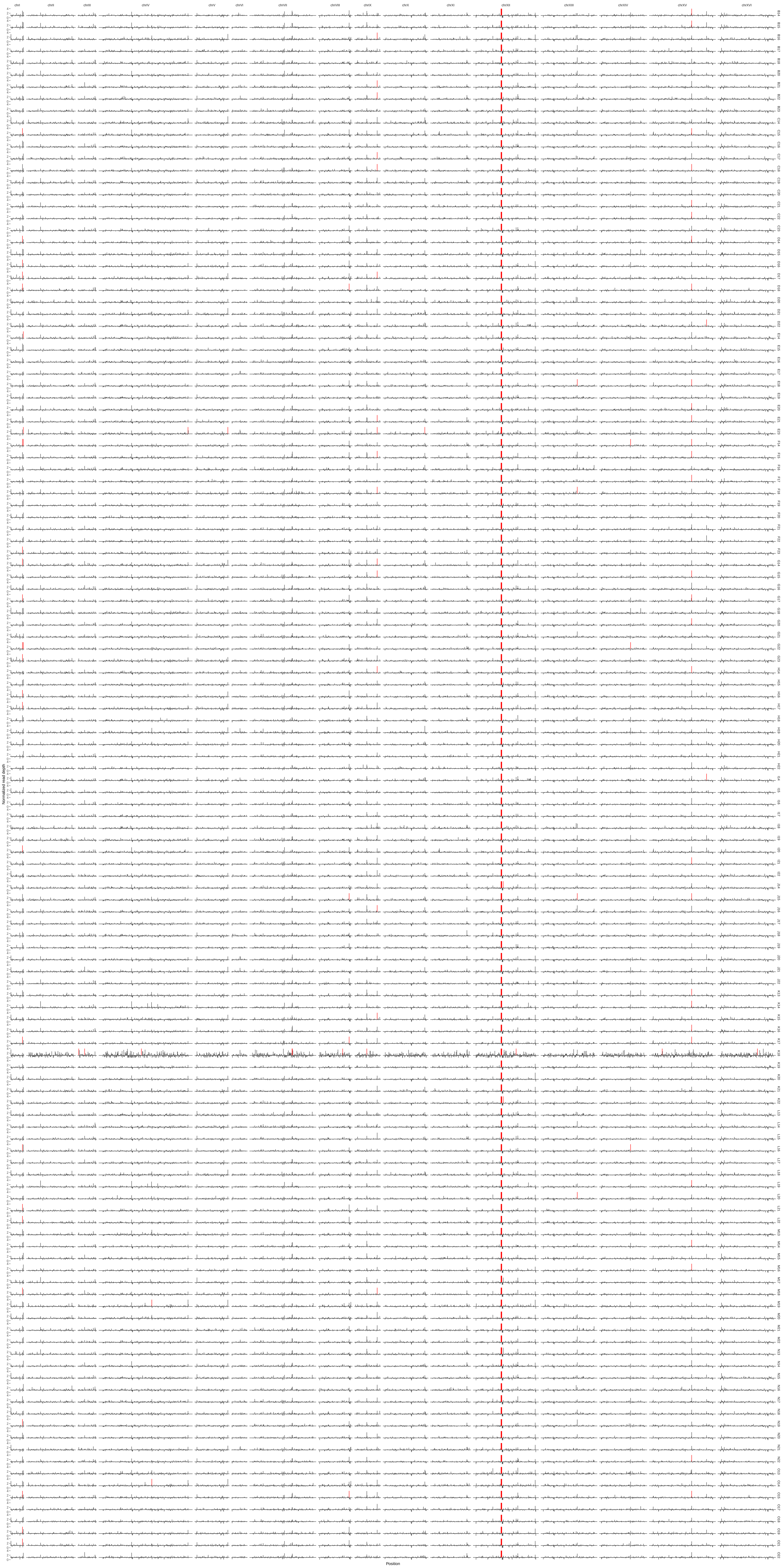
chrI
chrII
chrIII
chrIV
chrV
chrVI
chrVII
chrVIII
chrIX
chrX
chrXI
chrXII
chrXIII
chrXIV
chrXV
chrXVI
Normalized read depth
Position

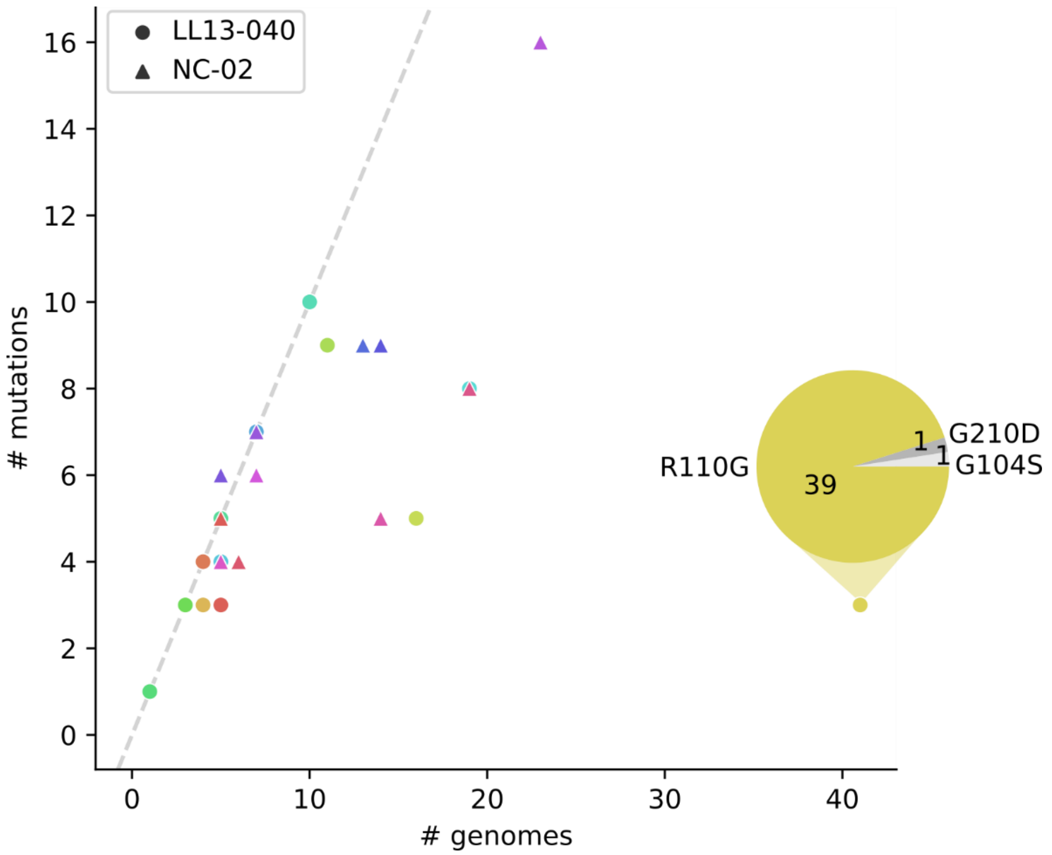


**Supp Fig 3. Diversity of Fur1 mutations per preculture.** Each marker (dot or triangle) represents a single preculture, from which mutants were selected (# genomes). A gray dashed line indicates if as many mutations have been identified in Fur1 as the number of genomes that carried them. For one outlier (only three mutations found in 41 strains which arose from the same preculture), a pie chart details the corresponding mutations and the number of strains that carried them.

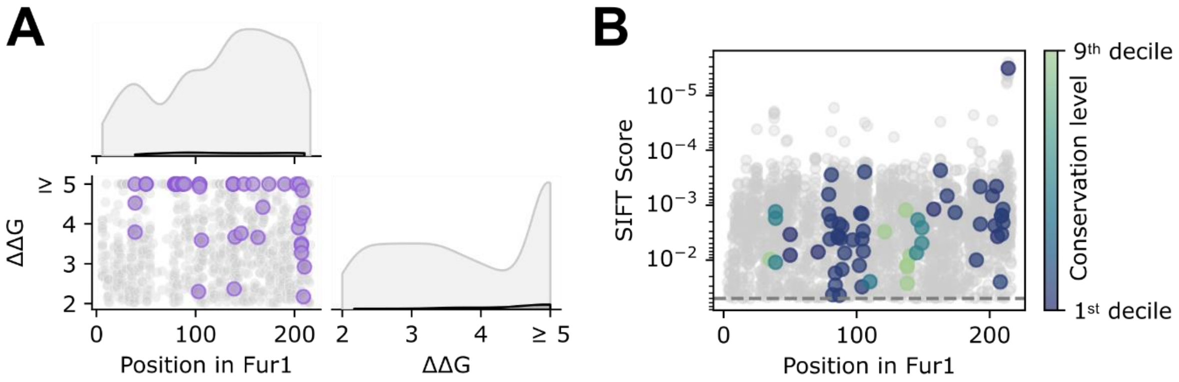


**Supp Fig 4. Predicted impact of Fur1 mutations.** Effect on stability (A) and conservation (B) of all possible amino acid substitutions in Fur1 which are predicted to be impactful by mutfunc [43]. Substitutions predicted to be non-deleterious are not shown (n=2,825 and n=1,437 out of 4,104 possible substitutions for stability and conservation, respectively). The substitutions captured in our dataset are highlighted (bigger colored dots, n=42 and n=67 out of 76 captured mutations for stability and conservation, respectively). (A) Side plots indicate the kernel densities for all data points (lightgrey) and substitutions captured in our dataset (black) along the position in Fur1 (top) or the ΔΔG value (right). (B) SIFT scores are represented on an inverted y-axis and are analogous to a *p*-value. Scores < 0.05 (gray dashed line) indicate a predicted deleterious mutation, with a low value (top of the plot) indicating that the amino acid change is very likely to affect protein function based on sequence conservation. For substitutions captured in our dataset, the color indicates the conservation level of the residue at that position.

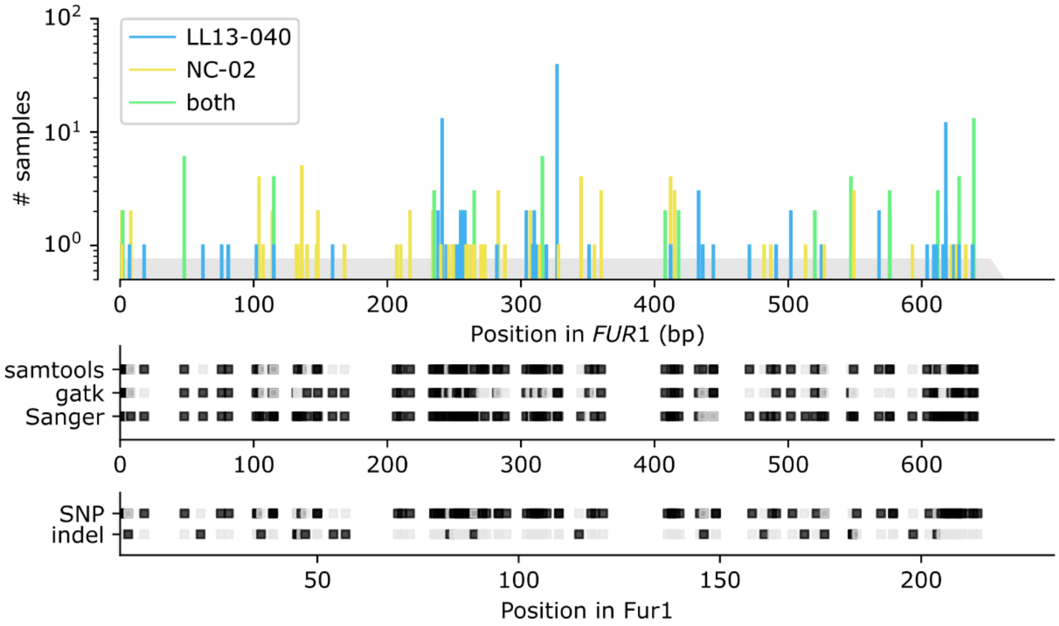


**Supp Fig 5. Location and type of mutations along the *FUR1* sequence.** The location of all detected mutations in the *FUR1* gene sequence (n=118, length of the gene represented by a gray half-arrow) is indicated by a barplot (first track), where every bar represents a unique mutation, and their height represents the number of unique genomes in which the mutation was identified. Bars are color-coded to indicate in which background the mutation was identified. The second and third tracks contain boolean indicators for the method of detection and the type of mutation (black for true, gray for false). On the third track, the corresponding positions along the Fur1 protein sequence are indicated.

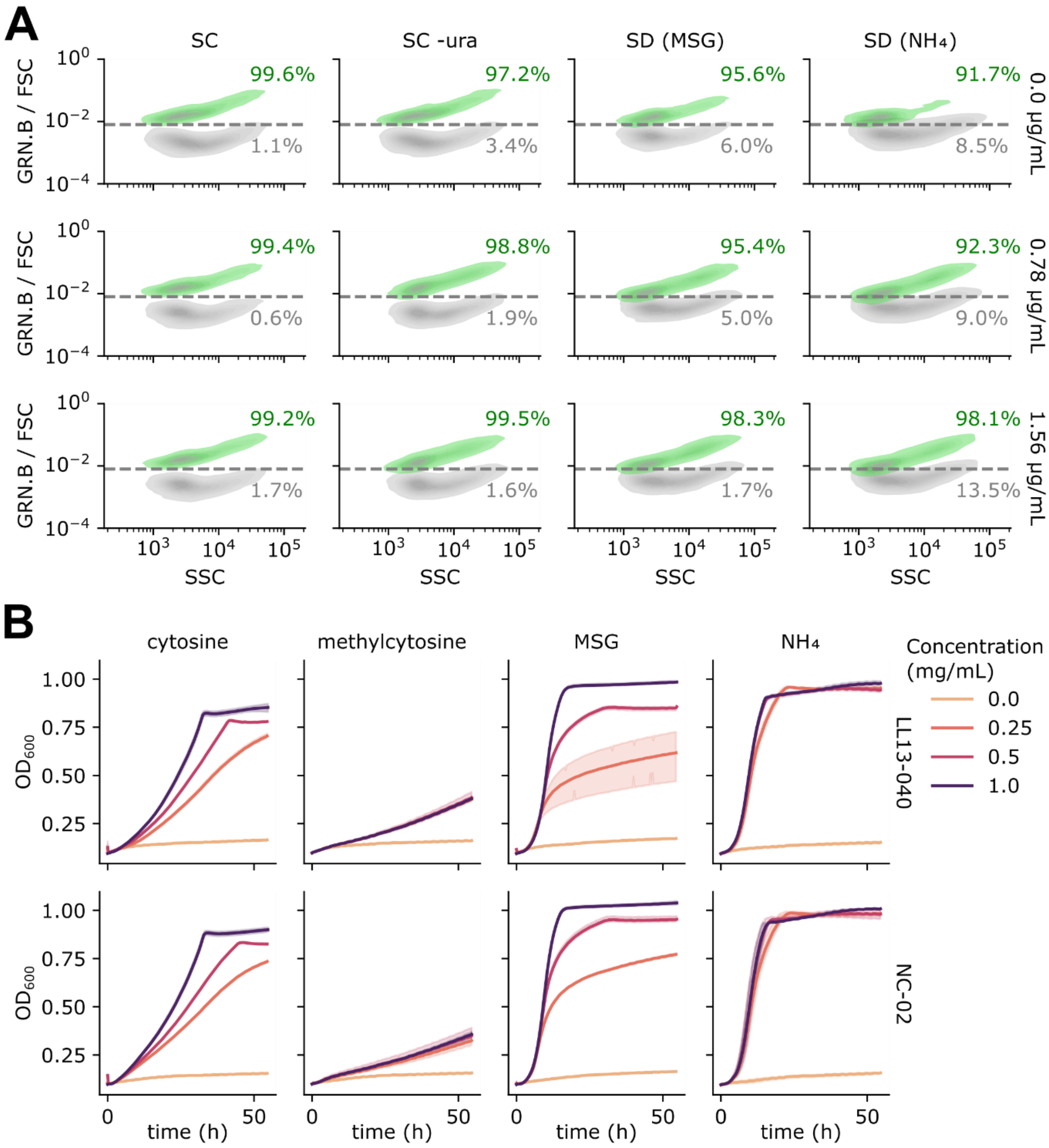


**Supp Fig 6. Fcy1 functionality in parental strains.** A) Density plots showing the fluorescence signal of *FCY1*-mEGFP (green), compared to the control without fluorescent reporter (gray) in the background LL13-040. Signal was acquired by cytometry for 5,000 events in the green channel (GRN.B) and normalized by the FSC value (proxy for cell size). Plots show the obtained value (GRN.B / FSC) as a function of the SSC value (proxy for cell granularity). A single threshold (gray dashed line) was used to indicate relative percentages of events for both strains in each condition. B) Growth was measured for both parental strains in YNB + 2% glucose containing different nitrogen sources to confirm *FCY1* functionality (deamination of the imported compound by Fcy1 leads to the obtention an intracellular nitrogen source).

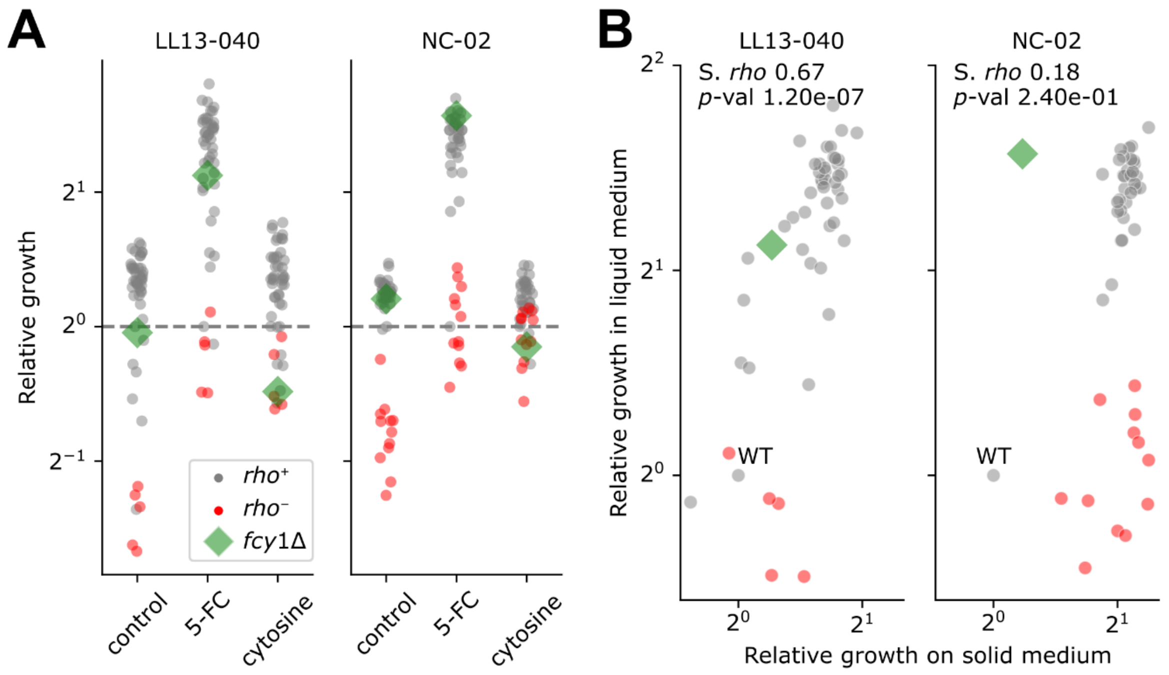


**Supp Fig 7. Fcy1-mediated resistance in liquid and solid media.** A) Growth assay in liquid medium. Cultures were inoculated from single replicates in a 384-well plate containing SD (control), SD + 25 µg/mL 5-FC (5-FC) or YNB + 2% glucose + 250 µg/mL cytosine (cytosine). Relative growth corresponds to the area under the curve (AUC, calculated on 13 h) normalized by the WT. B) Relative growth measured in liquid medium (data from panel A) compared to the one measured on solid medium for the corresponding mutants (data from Fig 2) at equal concentrations of 5-FC, with the corresponding Spearman's rank correlation coefficient (S. *rho*) and *p*-value.

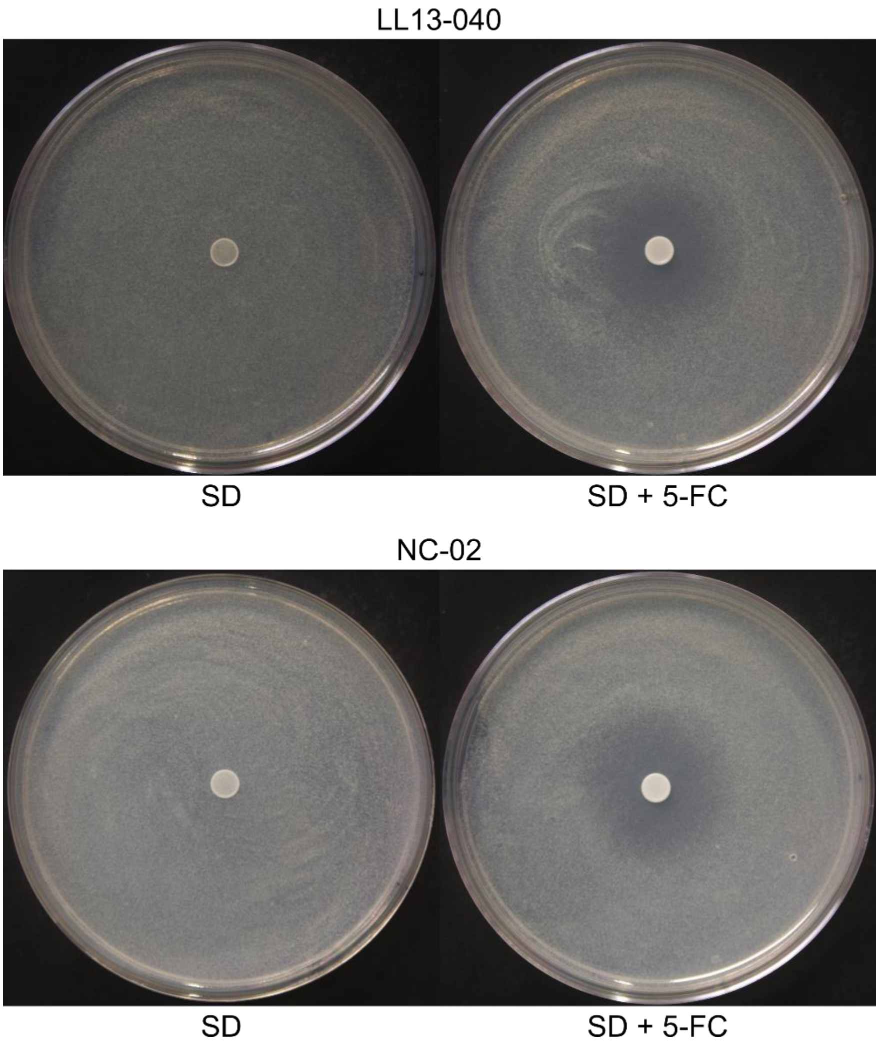


**Supp Fig 8. 5-FU cross-feeding inhibition zones.** Uncropped pictures as shown in Fig 6D.

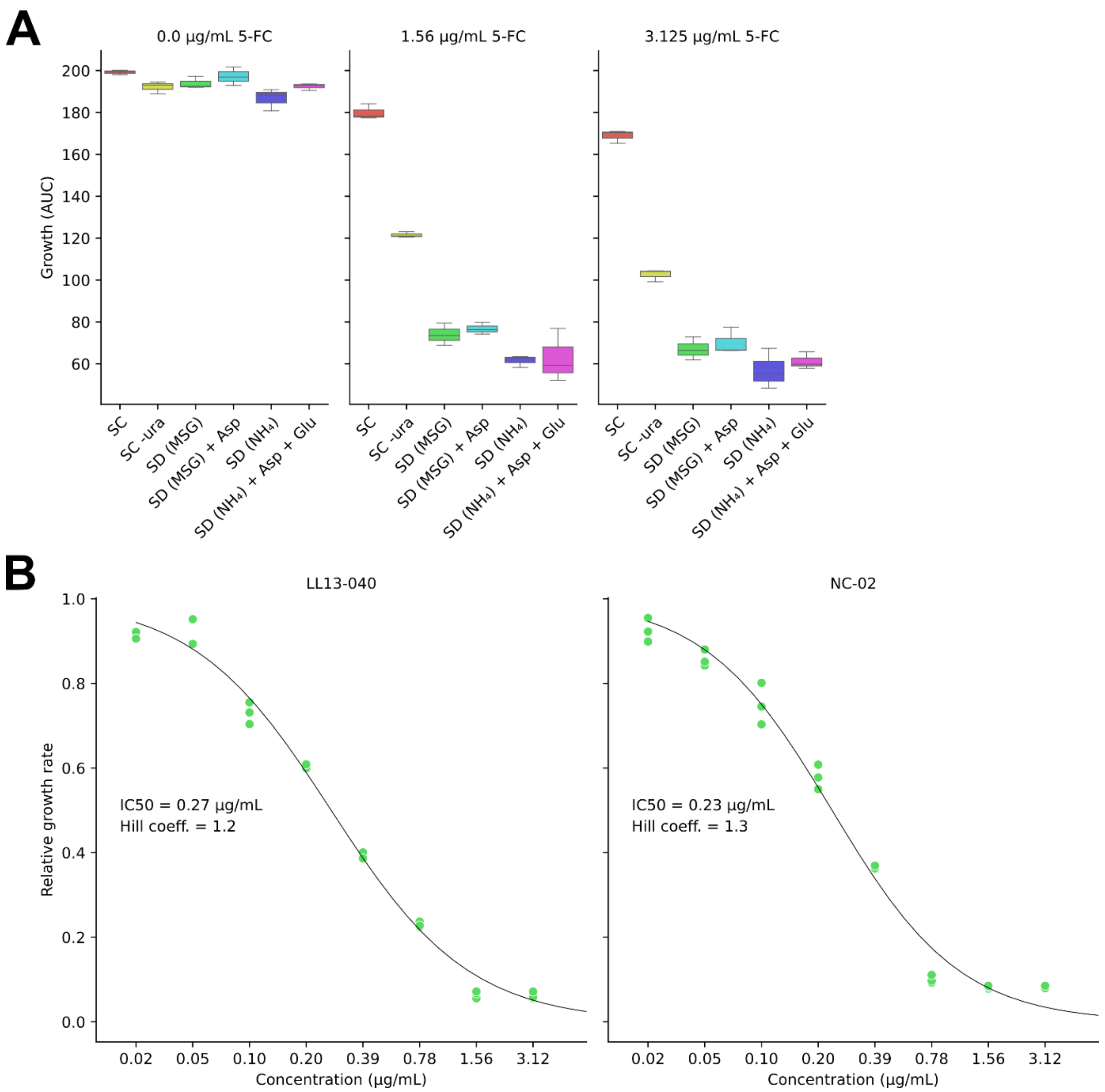


**Supp Fig 9. 5-FC dose response.** A) Growth assays for strain LL13-040 in different media supplemented or not with either 1.56 or 3.125 µg/mL 5-FC: synthetic complete medium with standard drop-out mix (SC complete), SC without uracil (SC -ura), SD (MSG) with or without 0.2% aspartate and SD ($NH_4$) with or without 0.2% aspartate / 0.2% glutamate. The area under the curve (AUC) parameter was calculated from three biological replicates. B) 5-FC dose-response curves in SD (MSG) for LL13-040 and NC-02. The concentration is shown on a log2 scale. 0 µg/mL 5-FC was used to normalize growth values. For each strain, the mean of three biological replicates for each concentration was used to fit the Hill equation. The corresponding IC50 and Hill coefficients are indicated.

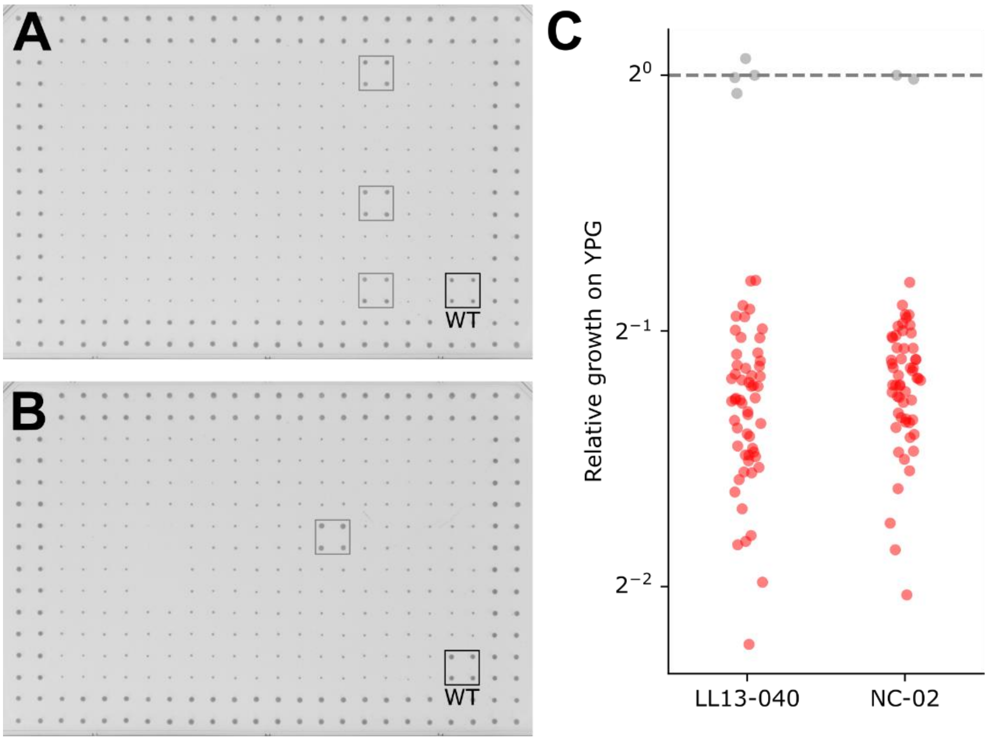


**Supp Fig 10. Growth measurements of colonies on YPG agar medium.** A, B) Pictures of arrays on YPG agar medium after 22 h incubation at 30°C for LL13-040 (A) and NC-02 (B). Each mutant is spotted in four replicates with the *fcy1*Δ mutant occupying the border as positive control. Grey squares highlight mutants initially misannotated as *rho*$^-$. Pictures were cropped and converted into inverted gray levels for clarity and downstream analysis of colony size. C) Growth curves were obtained by automatic detection of colony size on transformed pictures taken every 2 h for 22 h at 30°C. Relative growth corresponds to the mean area under the curve (AUC) normalized by the WT. *rho*$^-$ mutants included in all other figures are colored in red. Grey dots correspond either to the WT control (on the dotted line corresponding to a relative fitness of 1), or to the misannotated mutants mentioned above, which were therefore excluded from all analyses except the rhodamine accumulation experiment.
